# Genetic manipulation of Bacteriophage T4 utilizing the CRISPR-Cas13b system

**DOI:** 10.1101/2024.07.15.603621

**Authors:** Yuvaraj Bhoobalan-Chitty, Mathieu Stouf, Marianne De Paepe

## Abstract

CRISPR-Cas type II and type V systems are inefficient in modifying bacteriophage T4 genome, due to the hypermodification of its DNA. Here, we present a genome editing technique for bacteriophage T4 using the type VI CRISPR-Cas system. Using BzCas13b targeting of T4 phage, we were able to individually delete both T4 glucosyl transferase genes, *α-gt* and *β-gt*. Furthermore, we employed this method to mutate two conserved residues within the T4 DNA polymerase and to introduce the yellow fluorescent protein (YFP) coding sequence into T4 phage genome, enabling us to visualize phage infections. The T4 genome editing protocol was optimized to obtain recombinant phages within a 6-hour timescale. Finally, spacers homologous to a variety of T4 genes were used to study the generality of Cas13b targeting, revealing important variability in targeting efficiency. Overall, this study presents a detailed description of the rapid and easy production of T4 phage specific mutants.

**IMPORTANCE:** The use of phages for therapeutic purposes requires a complete understanding of their life cycle. For this purpose, it’s very useful to have a wide range of phage genome engineering tools at our disposal, each adapted to a particular phage or situation. Although T4 phage has been studied extensively over the past seven decades, a complete understanding of its lytic cycle is still lacking. Cas9- and Cpf1-dependent genome editing techniques for T4 have proven to be inconsistent due to the glucosyl-hydroxymethyl modification of the cytosine residues in its genome. The RNA targeting of the Cas13 system presents an ideal alternative, as demonstrated here, to overcome interference from DNA hypermodification. Apart from demonstrating a new genome editing technique in T4, we have generated a range of T4 variants demonstrating the efficacy of our technique in obtaining meaningful and difficult to construct phage mutants.

## INTRODUCTION

Since their discovery, the ability of Clustered Regularly Interspaced Short Palindromic Repeats and associated genes (CRISPR-Cas) systems to sequence-specifically cleave nucleic acids has been exploited for the genetic manipulation of organisms across all domains of life (1, 2). CRISPR-Cas are divided into Class 1 and Class 2 systems based on the complexity of the ribonucleoprotein interference complexes, and further divided into types I, III, IV (belonging to Class 1) and types II, V, VI (belonging to Class 2) (3). The type II CRISPR-Cas system Cas9, with its ability to introduce double-strand DNA breaks, is the most widely utilized CRISPR-Cas system in genome editing.

Bacteriophage T4, infecting *Escherichia coli*, is one of the best-studied bacteriophage. Historically, T4 mutants were generated based on induced (4, 5) or spontaneous (6) mutagenesis followed by selective pressure, such as temperature sensitivity or resistance to phosphonoacetic acid (7–9); all laborious, time-consuming and non-specific mutagenesis techniques. Specific gene modification techniques using homologous recombination with a template plasmid have also been largely used, but remained laborious to purify the recombinant in the absence of an easily selectable phenotype, and did not permit to obtain deleterious mutations (10, 11). Over the past decade, phage genome editing techniques utilizing homologous recombination coupled with selection of recombinants by CRISPR-Cas targeting have been developed for *E. coli* bacteriophages T3, T4, T5, T7, lambda, P1 and *Klebsiella* phages (11–17). However, these techniques rely on double-strand DNA targeting dependent CRISPR-Cas systems, Cas9 and Cpf1. CRISPR-Cas type II and type V dependent genome editing of T-even phages has been shown irregular due to hypermodification of the cytosine residues in the phage DNA (13, 16, 18–22). Hypermodification of cytosines in T4 was previously demonstrated to be important for distinguishing between host and phage genetic material, enabling protection of the phage against restriction-modification systems (23, 24) and T4 driven host genome degradation, and implementing transcriptional specificity (25).

Cas13, belonging to the type VI CRISPR-Cas system, is unique in that while retaining the sequence specificity of class 2 CRISPR-Cas systems, it recognizes RNA targets instead of DNA like the types II and V CRISPR-Cas systems (26, 27). Apart from the cleavage of the target RNA based on sequence specificity, it also contains a non-specific trans-RNA collateral cleavage activity, regulated by the protospacer containing target RNA (27, 28). To date, type VI CRISPR-Cas system has been further classified into four Cas13 phylogenetic subtypes, from Cas13a to Cas13d, presenting limited amino-acid sequence similarity. RNA targeting by CRISPR-Cas13 system is dependent on the crRNA efficiency (29), expression level of the target transcript (30) and protospacer-flanking site (PFS) requirements (31–33). Recently, the Cas13a system was repurposed for genome editing in phage T4 (34) and *Pseudomonas aeruginosa* nucleus forming Jumbo phage ΦKZ (29), whose DNA is protected from nucleases. As CRISPR-Cas13 subtypes present differences in binding and cleavage specificity, we intended to use the diversity among CRISPR-Cas13 system to expand the toolbox for genome editing.

Here, we utilize the *Bergeyella zoohelcum* Cas13b system to establish a genetic manipulation technique for phage T4. Using a dual host system, first to introduce the desired edit and then latter to select the recombinant phage, we were able to perform gene deletions (T4 glucosyl transferases), point mutations (within *gp43*, the T4 DNA polymerase gene) and gene insertion (of Yellow Fluorescent Protein gene *yfp*). Notably, we could obtain a highly deleterious mutation, thanks to efficient counterselection of wild-type phage. The protocol was optimized to obtain edited phages within 6 hours following the initial infection. We present here an encompassing view of Cas13b based genome editing, including its drawbacks, as well as the new isogenic T4 mutants obtained with this technique.

## RESULTS

### Unmarked deletion of T4 *alpha*- and *beta*-*glucosyltransferases*

In contrast to the DNA-dependent targeting characteristic of the Class 2 CRISPR-Cas types II (Cas9) and V (Cpf1), the type VI (Cas13) system recognizes and cleaves RNA. Despite requiring target transcription, we reasoned that the Cas13 effector activity would be more uniform across different targets on T4 due to its non-dependence on base modifications of the transcribing protospacer containing DNA.

Previously, it had been shown that among the classified Cas13 systems, Cas13a and Cas13b were the most widespread subtypes, with the former dispersed among several bacterial phyla (34, 35). Accordingly, we chose to work with Cas13a (C2c2) from *Leptotrichia shahii* (27) and Cas13b from *Bergeyella zoohelcum* ATCC 43767 (31). First, we compared the targeting efficiency of Cas13a and Cas13b against the T4 genes *alpha-glucosyltransferase* (*α-gt*) and *beta-glucosyltransferase* (*β-gt*). Cas13 subtype specific protospacers were identified and corresponding spacers were cloned into the respective plasmids pC0003 (expressing Cas13a) and pBzCas13b. A 6-log decrease in T4 plaquing ability was observed upon expression of either *α-gt* or *β-gt* specific spacers along with Cas13b as compared to Cas13a, which had no significant impact on the plaquing ability (Figure 1A). Based on these results we concluded that targeting of T4 by BzCas13b was likely to be also more effective on others targets and hence all subsequent experiments were conducted utilizing this subtype.

**Figure 1.**
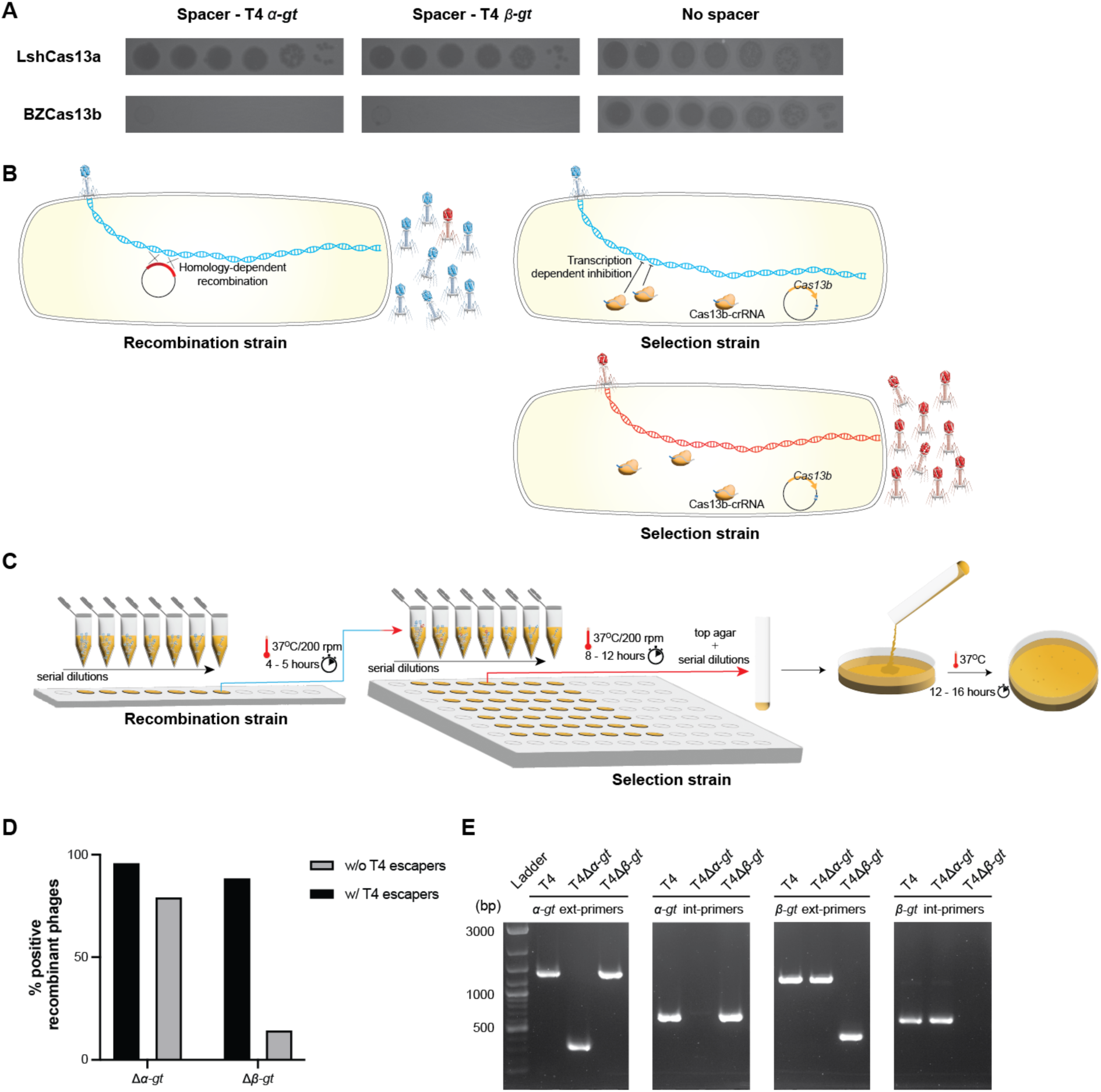
Cas13b based deletion of T4 alpha- and beta-glucosyltransferase genes. **A.** Spot assay of 10-fold serial T4 dilutions, comparing its infectivity in *E. coli* strains expressing either plasmid-borne *Leptotrichia shahii* Cas13a (Top panel) or *Bergeyella zoohelcum* Cas13b (bottom panel), in the presence of corresponding spacers targeting either T4 *α-gt* and *β-gt*, or in the absence of spacer. **B.** Illustration of the two-step genome editing method used. In a first step, a recombination strain carrying a plasmid containing the homologous arms necessary for deletion of the target gene was used to propagate wild-type T4. Subsequently, the resulting phage lysates were used to infect a selection strain, carrying a CRISPR-Cas13b plasmid along with a spacer specific to the transcript of the target gene, to purify the recombinants. **C.** Illustration of the practical details of the methodology. Both phage amplification steps were realized in 200 microliter cultures, at different M.O.I. Cultures that showed clear signs of lysis in the first step were serially diluted and used to infect the selection strain. **D.** Percentage of wells with recombinant phages following infection of the selection strain, relative to the total number of wells that showed clear signs of lysis. The percentages of wells comprising escaper variants (w/ T4 escapers) and those with only the expected recombinants (w/o T4 escapers) are shown. The respective PCR confirmations are shown in Supplementary Figure 2 and Supplementary Figure 3. **E.** PCR confirmation of the purified recombinant phages T4Δ*α-gt* and T4Δ*β-gt*, using external primers, across the *α-gt* (*α-gt* ext primers) or *β-gt* (*β-gt* ext primers) and internal primers, within the *α-gt* (*α-gt* int primers) or *β-gt* (*β-gt* int primers). Ladder : 1 kb+ DNA ladder.

Type II and V CRISPR-Cas targeting introduces a dsDNA break into the target genome, which improves the homologous recombination-repair process and simultaneously selects for the recombinant phage. RNA targeting by Cas13 does not introduce any nicks in the genome and is inherently dependent on the systems encoded within the host or the phage to promote recombination, independent of any induced dsDNA breaks. To overcome this disadvantage, we adopted a two-step system (11, 34, 36), the first step to introduce the desired modification and the second to select for the recombinant phage, as illustrated in Figure 1B.

For the deletion of both *α-gt* and *β-gt*, homologous arms (approx. 80bp in length) identical to the respective 5’ end and 3’ end of the genes were fused together (forming the desired deletion) and cloned into the pBAD24 vector. *E. coli* DH10B strain carrying the resulting plasmid was infected at different MOI (multiplicity of infection) of wild-type T4 and incubated at 37°C for 5 hours (Figure 1C and Supplementary Figure 1). Subsequently, the supernatants of each culture, containing a mixture of recombinant and wild-type T4, were serially diluted and propagated in the selection strain, carrying the plasmid-borne Cas13b and a spacer targeting either *α-gt* or *β-gt*, at 37°C for 8-12 hours (Figure 1C and Supplementary Figure 2). The cultures that showed cell lysis were then screened by PCR for the presence of recombinant phage. In case of *α-gt* deletion, among the cultures that showed lysis (corresponding to MOI above 10^-3^), most contained only the recombinant phage and a small minority also contained phage escapers (Figure 1D and Supplementary Figure 2). In case of *β-gt* deletion, the same pattern was observed, though with a larger proportion of cultures containing also wild-type phage or phage escapers in addition to the phage recombinants (Figure 1D and Supplementary Figure 3). In both cases, the residual amount of wild-type phage increased with the initial MOI. Single plaques of the recombinant phages (named T4Δ*α-gt* and T4Δ*β-gt*) were isolated and the deletion verified by PCR (Figure 1E).

Together, these results show that Cas13b from *Bergeyella zoohelcum*, in the presence of a suitable spacer, can efficiently inhibit the propagation of phage T4. Furthermore, this inhibitory effect can be harnessed to modify the genome of the T4 phage, and was utilized here to construct two new T4 phage variants, T4Δ*α-gt* and T4Δ*β-gt*. Sequencing of T4Δ*α-gt* and T4Δ*β-gt* genomes revealed they were isogenic to the ancestral phage, with the exception of a single point mutation in former and two point mutations in the latter (Supplementary Figure 4), demonstrating the specificity of the genome modifications introduced.

### Substitutions in the exonuclease domain of T4 DNA polymerase

To further demonstrate the suitability of the type VI CRISPR-Cas system for the genetic manipulation of bacteriophages, we introduced mutations into a functionally significant phage protein domain. T4 DNA replication is ensured by T4 DNA polymerase (T4 DNA pol), possessing a N-terminal proofreading exonuclease domain. Various mutants of the T4 DNA pol have been characterized, predominantly *in vitro*, including mutants in the exonuclease domain. Two amino acid residues in particular, Y320 and D324, have been shown to participate in the active site and metal ion coordination, respectively. Reduced polymerase and exonuclease activities were observed upon the substitution of amino acids Y320 and D324 by alanine, compared with the activities of the T4 wild-type polymerase enzyme (37). To introduce these mutations, a DNA fragment of approximately 160 bps, spanning the site to be modified with nucleotide mutations necessary for the substitution of Y320 and D324 by alanine, was cloned into a plasmid vector (recombination plasmid). Additional silent mutations were also introduced into this recombination fragment to aid the recombinant phage escape targeting by Cas13b (Figure 2A) (38, 39). Genome editing was performed as described previously; first by multiplying T4 on a strain carrying the recombination plasmid containing the polymerase gene fragment to be modified, and then by growing the mixture of wild-type and recombinant phages on a selection strain expressing Cas13b and a spacer targeting the wild-type DNA pol (Supplementary Table 1). The mutations within the T4 DNA polymerase recombinant phage (named T4DNApolMut01) was confirmed both by PCR (Figure 2B) and Sanger sequencing (Supplementary Figure 5).

**Figure 2.**
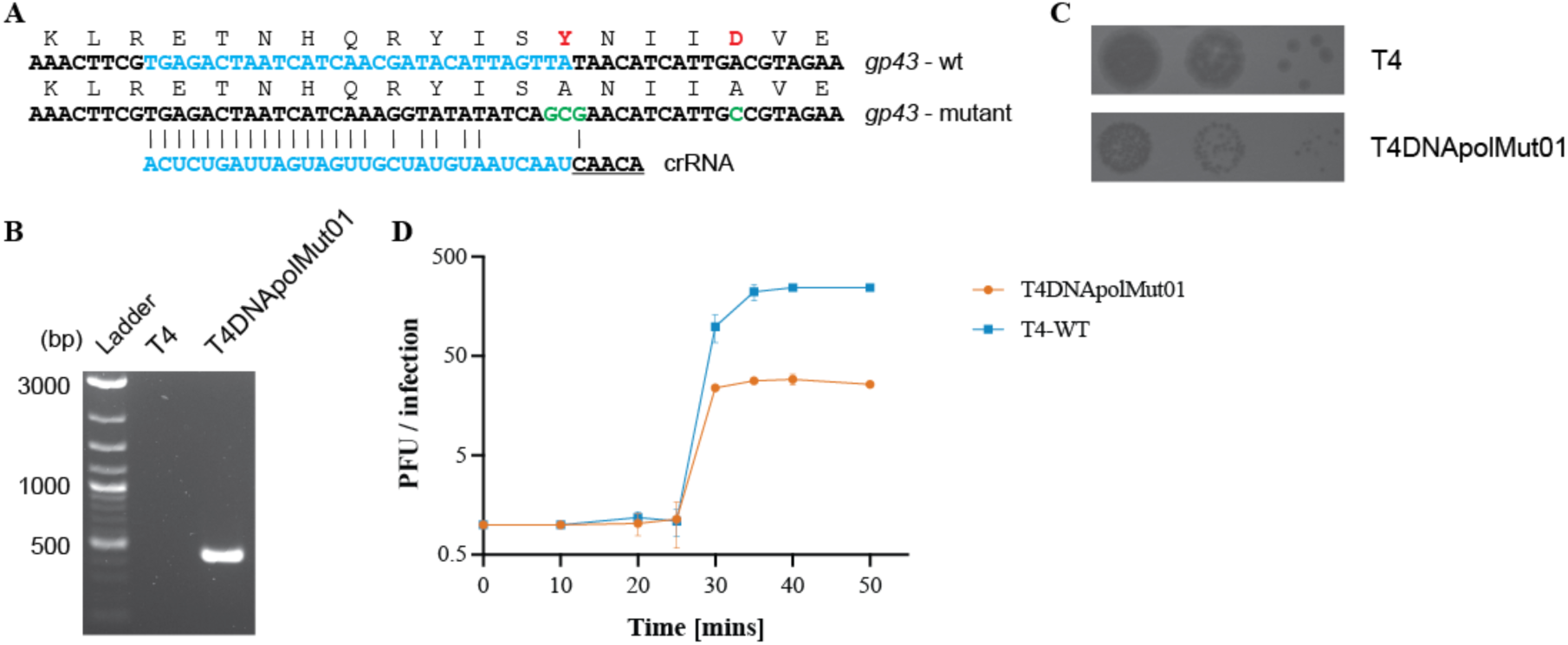
Mutation of residues in the exonuclease domain of T4 DNA polymerase. **A.** Amino acid and nucleotide changes in *gp43* mutated gene (*gp43_Y320AD324_*) compared to wild- type gene, and sequence of the spacer targeting only the wild-type gp43 sequence. Additional mutations were introduced to improve mutant phage targeting escape. **B.** PCR confirmation of the presence of the desired mutations in T4DNApolMut01, with the wild-type T4 phage used as a negative PCR control. Primer pair: T4_DNApol_chk_for and DNApolY320Amut_chk_rev. The reverse primer has a reduced binding affinity for the wild-type *gp43* gene. **C.** Spot assay of the wild-type T4 phage and the mutant phage showing differences in their plaque size. **D.** One-step growth curve of wild-type T4 and T4DNApolMut01on *E. coli* DH10B. Data shown are the mean of three biological replicates, represented as mean value ± SD.

In comparison to the wild-type T4 phage, the plaque size of T4DNApolMut01 was considerably smaller (Figure 2C). We estimated the burst sizes of wild-type T4 and T4DNApolMut01 to be 245 and 29, respectively, which corresponds to a 8.5-fold decrease associated with the DNA Pol mutations (Figure 2D). Despite the mutations in the exonuclease domain affecting the DNA synthesis rate and subsequently the virus yield, the latency period of the virus remained unaffected, with both wild-type and mutant phage lysing *E. coli* cells within a window of 25 to 30 minutes post infection. This mutation is therefore highly deleterious for the phage, disfavoring the growth of recombinants.

Taken together, these results show that Cas13b, with significant number of mismatches in the protospacer region, can be exploited to introduce point mutations into T4 genome, even deleterious ones, further demonstrating that the type VI system can be used as a tool for genome editing in bacteriophages.

### Tracking T4 infection in single-cells

Taking advantage of the editing technique established, we aimed to visualize phage infection by expressing a fluorescent protein under the control of a native T4 phage promoter. Since we aimed to visualize infection from the beginning of the lytic cycle, we choose to insert the fluorescent protein gene in T4 *β-gt*. Indeed, T4 *β-gt* is a T4 early gene, expressed within minutes after infection (Supplementary Figure 6), from the promoter of the adjacent dCMP hydroxymethylase *gp42* gene (40). For the same reason, because of its very short maturation time (*t*_50_ at 37°C), we reasoned that mVenusNB (Yellow-green), with a *t*_50_ value of 4.1 mins, would be an appropriate fluorescent protein (41). The *mVenusNB* coding sequence was thus inserted in between 80-100 bps long arms, homologous to the upstream and downstream intergenic regions flanking the *β-gt* gene in T4 (Figure 3A), and cloned into the recombination plasmid. Additionally, the native *β-gt* SD (Shine-Dalgarno) sequence was modified into a stronger SD sequence (Supplementary Figure 7). Like before, the *E. coli* DH10B strain carrying the recombination plasmid was infected with wild-type T4 phage, following which the recombinant phages were isolated using a selection strain expressing spacers targeting *β-gt*. T4 recombinants containing *mVenusNB* coding sequence, named T4Δ*β-gt*∇*mVenus,* was confirmed by PCR on individual plaques (Figure 3B). The insertion was estimated to slightly reduce the burst size of the T4Δ*β-gt*∇*mVenus* phage compared with wild-type T4 phage (Figure 3C and Figure 2D).

**Figure 3.**
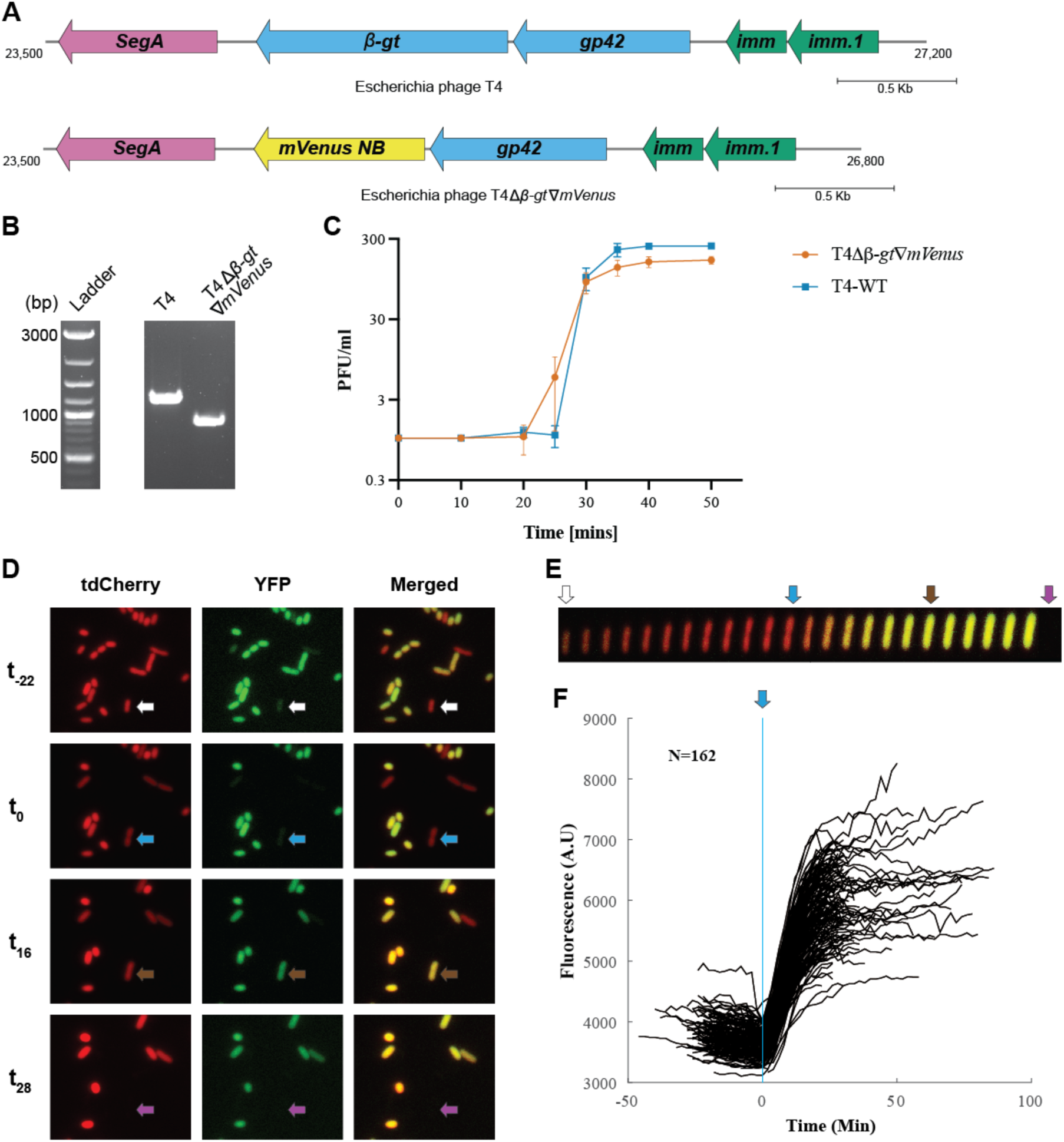
Construction and analysis of *yfp* encoding T4 phage. **A.** Gene map of *β-gt* neighborhood in wild-type T4 (upper panel) and in the intended recombinant phage with the *yfp* replacing the *β-gt* gene. **B.** PCR confirmation of the replacement of *β-gt* with *mVenusNB*, the wild-type T4 phage was used in the control reaction. **C.** One-step growth curve of T4Δ*β-gt*∇*mVenus*. Data shown are mean of three biological replicates, represented as mean value ± SD. **D.** Fluorescent microscopy images of *E. coli* MG1655 PRNA1::*tdCherry* upon infection with T4Δ*β-gt*∇*mVenus* at four different times. From top to bottom: -22, 0, 16 and 28 minutes from start of infection of the cell identified by the arrow, with tdCherry, YFP and merged channels from left to right. Image acquisition on agar pads started approximately 20 minutes after the mixing of phage and bacteria (<u>Supplementary Data 1</u>). **E.** Kymograph of the single cell identified in D, with tdCherry and YFP merged signals. **F.** YFP mean fluorescent intensity evolution with time of 162 infected cells, synchronized based on the time point at which elongation stops.

To characterize T4Δ*β-gt*∇*mVenus* phage derived yellow fluorescence during infection, we tracked by fluorescence microscopy infected *E. coli* MG1655 *PRNA1::tdCherry* cells (Supplementary Table 3). All *E. coli* cells express red fluorescence from the tdCherry protein, while T4 infected cells were identified by yellow fluorescence (Figure 3D, Figure 3E and <u>Supplementary Data 1</u>). Besides the appearance of yellow fluorescence, infection is also characterized by an abrupt decrease of cell elongation and an increase in cell width (Figure 3E) enabling to estimate the time of infection. Most infected cells lyse 25 to 30 minutes after cell elongation has stopped, which corresponds to the duration of T4 life cycle as determined from the single burst assay, corroborating that elongation stop corresponds to the time of infection (Figure 3C). Some cells lyse only after 90 minutes, which is expected due to the lysis inhibition phenotype expressed by T4 phage. Analysis of fluorescence intensity in 162 lysing cells, synchronized based on the time at which cell elongation stops, demonstrates a sharp increase in fluorescence intensity few minutes after infection, until reaching a plateau towards the end of the phage life cycle (Figure 3F), in line with previously described β-gt protein expression (40).

Overall, we have demonstrated the possibility of gene insertion in T4 genome thanks to Cas13b targeting, and shown the interest of such construction towards tracking T4 infection in single cells.

### Optimizing T4 genome editing

Although the previously described Cas13 based genome editing protocol is already efficient, we aimed at reducing the time necessary to obtain phage mutants by shortening both the recombination and the selection steps. A way to shorten the recombination step is to infect cells at a high MOI, as we have shown previously that despite decreasing the number of phage lytic cycles on the recombinant strain, and therefore the chances of recombination, a high MOI still permitted us to obtain phage mutants (Supplementary Figure 2 and Supplementary Figure 3).

In order to test a shortened procedure, the recombination strain for the deletion of *α-gt* was infected at an MOI of 0.1 and 1.0 and incubated at 37°C for only 60 minutes (instead of 5 hours as demonstrated previously), which was sufficient to ensure complete host lysis. Subsequently, serial dilutions of the supernatants were directly spotted on the selection strain, carrying the plasmid-borne Cas13b and a spacer targeting *α-gt*, without first growing the mixture on the selection strain. Plates were incubated at 37°C for five hours (Figure 4A). Among the 32 individual plaques screened by PCR, 30 corresponded to the recombinant phage (Figure 4B and Supplementary Figure 8). This quick protocol was repeated for the deletion of *β-gt*, also with a positive outcome, as 50% of screened plaques corresponded to the recombinant phage.

**Figure 4.**
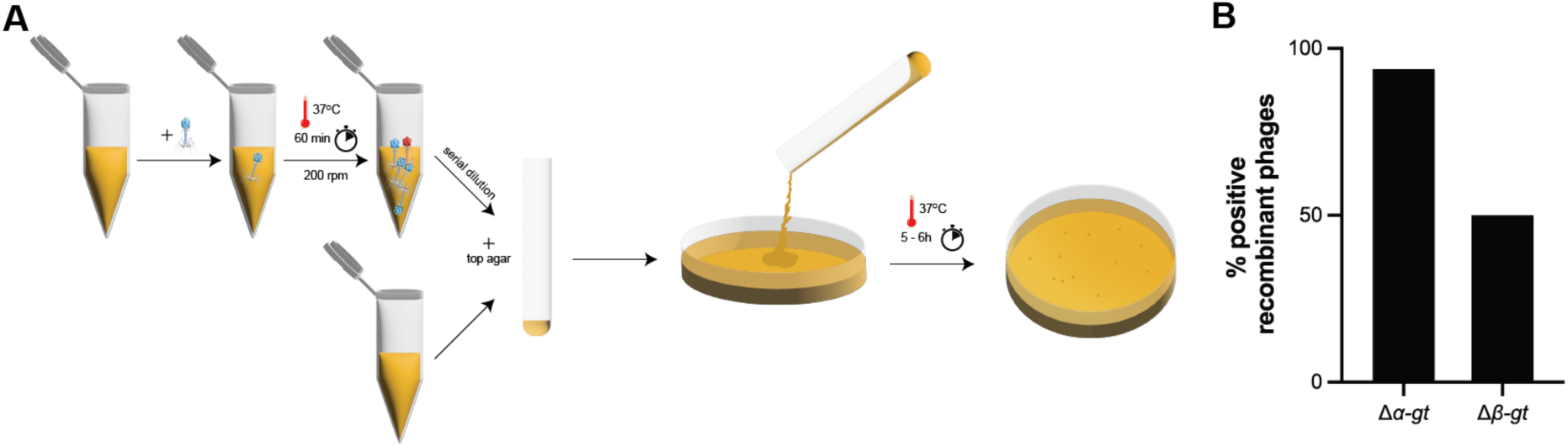
Fast T4 genome editing protocol. **A.** Illustration of the fast genome editing method. Wild-type T4 phage is propagated on the recombination strain carrying the plasmid necessary for the homologous recombination dependent modification of the target, for 1 hour at 37°C. Subsequently, the phage lysate obtained was serially diluted, and directly plated on the selection strain, carrying a CRISPR-Cas13b plasmid along with a spacer specific to the transcript of the target gene, which blocks the propagation of the wild-type phage and enable to isolate the recombinant phage. **B.** Percentage of recombinant phages post selection with the selection strain, as shown by PCR (shown in Supplementary Figure 8).

Upon Sanger sequencing of the escaper phages across the protospacer region, we observed either large microhomology based deletions of the protospacer region or modifications within the protospacer region, preventing Cas13b-dependent targeting (Supplementary Figure 9). Overall, here we tested a quick protocol to obtain different types of mutations in phage T4, establishing the possibility of genome editing within 6 hours.

### Variability of Cas13b crRNA efficacy in T4

In order to better delineate the Cas13b targeting efficiency against different T4 genes, we introduced in the Cas13b encoding plasmid 29 spacers complementary to protospacers situated on 23 different T4 genes (Supplementary Table 1). As efficacy could depend on the timing and strength of transcription, we choose early genes (expressed during the first five minutes following infection), middle genes (expressed between 5 and 10 minutes) and late genes (expressed after 10 minutes). The protospacers were chosen based on the likelihood that the flanking sequence (PFS - Protospacer Flanking Sequences) would be suited for targeting by the Cas13b system (32), NAA sequence downstream of the protospacer along with maximal non-complementarity between the anti-tag and 5’-tag sequence of the spacer to reduce any anti-tag based inhibition of targeting (31, 42). Few spacers that do not follow these rules were also tested. As seen through spot assay of serial T4 dilutions, targeting by Cas13b varied from highly efficient to no efficiency (Figure 5). Some spacers resulted in a more than 5-log decrease in plaquing efficiency (*denA*, *gp43*, *dexA*, *gp7*, *gp10*), others had intermediate level of targeting, with 1-log to 4-log decrease in plaque formation (*alc*, *gp56*, *res*, *gp61*, *gp6* (x2), *Vs. 6*, *gp15*, *gp10*, *hoc*, *rIIA*, *gp18*, *gp12* (x2) and *uvsW*), and finally some spacers showed no targeting at all (*gp43*, *res*, *gp3*, *gp34*, *gp18*, *gp42*, *dda*, *gp30.3*, *gp23*) (Figure 5).

**Figure 5.**
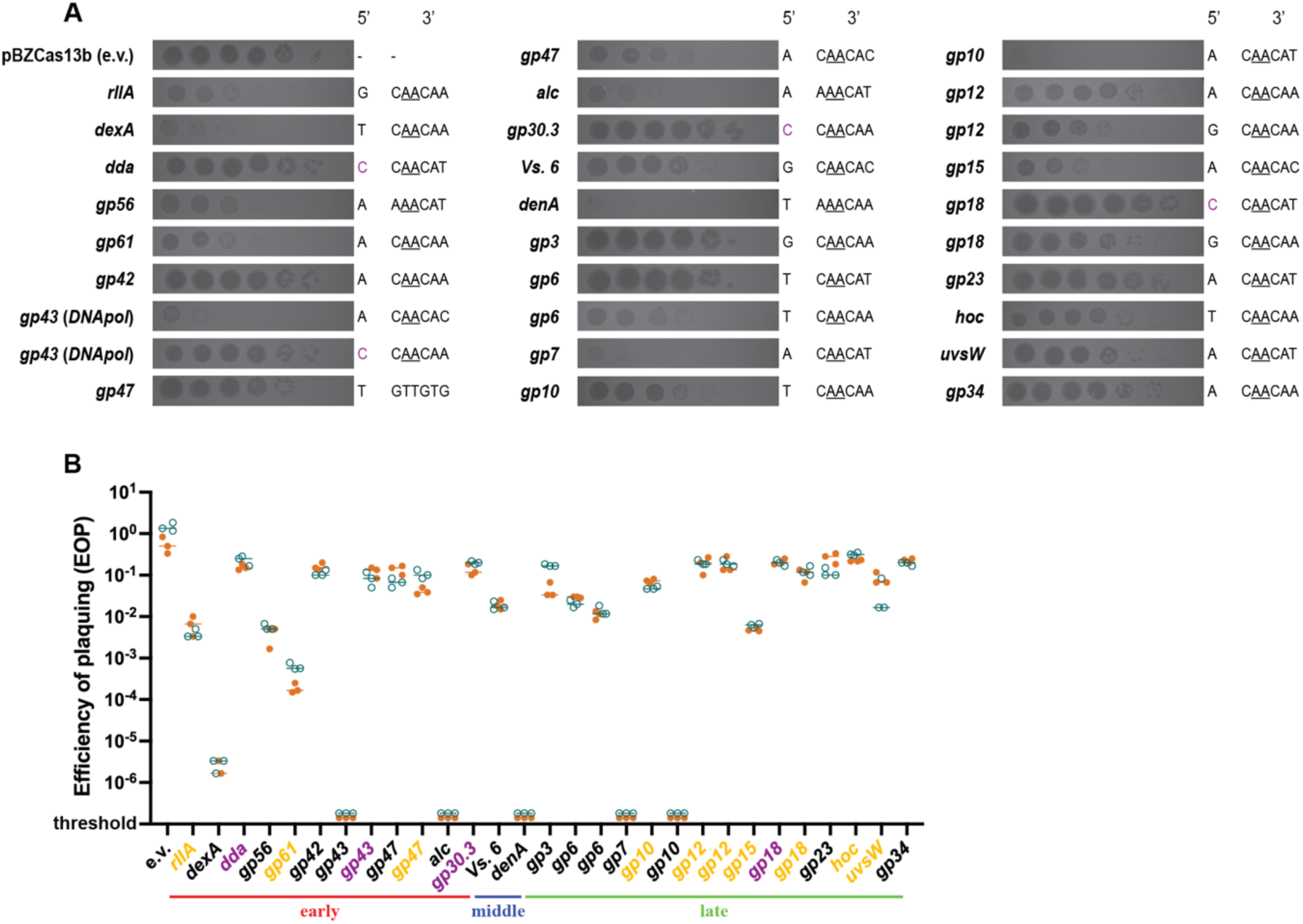
Plaquing efficiency with CRISPR-Cas13b spacers targeting different genes in phage T4. **A.** Dilutions of wild-type T4 were spotted on *E. coli* DH10B carrying Cas13b specific spacers inserted within the pBzCas13b plasmid. Gene names of the corresponding transcripts and the 5′- and 3’-protospacer flanking sequence (PFS) of the spacers are shown on either side of the respective spot assay image. Previously validated 3’ PFS motif (31), requiring adenine at positions -2 and -3 are shown (underlined) in all protospacers containing this motif. **B.** Quantification of the plaquing efficiency of T4 phage on different Cas13b spacer carrying strains of DH10B. Values shown are technical triplicates of two biological duplicates (shown in orange and deep teal). In some cases, the number of plaque forming units could not be estimated (shown as below threshold values). The target T4 genes (shown in X axis) are grouped according to their expression timing during lytic cycle into early, middle and late genes. The classification was based on a previous transcriptomic study (40). Genes highlighted in yellow represent Cas13b spacer targets resulting in significantly smaller plaques than in the absence of targeting, as shown in (A), regardless of the EOP value. Genes highlighted in purple represent protospacers with 5’ PFS of cytosine.

Among the 29 protospacers analyzed 28 contained the recommended 3’PFS and 24 protospacers followed both the 5’ and 3’ PFS recommendations (31). Despite this targeting was not uniform across all these 24 target mRNA. No clear pattern of transcription could be related to targeting efficiency. In addition, protospacer sequence-dependent differences in Cas13b- targeting were observed among single transcripts of *gp43*, *res*, *gp6*, *gp10*, *gp18* and *gp12*, reinforcing the idea that the difference do not (only) depend on the gene transcription pattern.

## DISCUSSION

DNA hypermodification is a frequent feature in strictly lytic phages (43), a phenomenon that most likely results from the coevolutionary arm race between phages and bacteria, as it enables phages to escape most restriction enzymes. DNA hypermodification also constrains the cross-target consistency of type II and type V CRISPR-Cas activity, in turn influencing the efficiency of genome editing, in hypermodified strictly lytic phages (18, 21, 22). Here, we report a CRISPR-Cas type VI-B based tool that can be utilized in the counter-selection of recombinant phage post genome editing, recombinants being obtained thanks to the inherent phage recombination machinery. In addition to the recent studies on bacteriophage genome editing using the type VI-A system, this study presents a crucial extension to the hypermodified phage genome editing toolbox.

Thanks to this technique, we obtained several T4 mutants of interest (Figure 1). In the mid 1960s, glucosyltransferase mutants of T2, T4 and T6 phages (referred to as T2gt, T4gt and T6gt) had been obtained based on non-specific chemical mutagenesis techniques and isolated as “host-defective” mutants (44, 45), and these mutants are still used today to study the impact of cytosine glucosylation on types II and V CRISPR-Cas system targeting (20, 21). The introduction of mutations outside the region of interest, as well as the possibility of reversion, are two main drawbacks of this classical mutation method, that might blur some results. The precise deletions of entire open reading frames obtained in our work guarantee that the phenotype observed cannot be related to other unintended mutations. In addition, through the insertion of a YFP fluorescent reporter within T4 genome, we were able to visualize individual infections by phage T4 (Figure 3). Applications include an accurate analysis of phage T4 life cycle in different conditions, as well as the determination of the expression pattern of individual T4 genes, providing a high-resolution alternative to RNA sequencing. Finally, introduction of point mutations that diminish the polymerase activity of the T4 DNA Pol demonstrates the possibility to obtain highly deleterious mutations using the BzCas13b system (<u>Figure2</u>).

The two previously published studies on T4 Cas13a targeting have reported contrasting results concerning the variability of efficiency across different protospacer regions (29, 34). The targeting of phage T4 using LbuCas13a was shown to have near similar efficiencies across different protospacers, despite them being located on phage transcripts with differing expression levels and patterns. In contrast, considerable variability in the targeting of phiKZ using LseCas13a was observed, both with protospacers corresponding to different transcripts and within the same transcript. The former study also reported lower efficiency and variability using the RfxCas13d system. In our study, we observed a 6-fold lower targeting efficiency with the LshCas13a system in comparison with the BzCas13b, with two different spacers targeting the *α-gt* and *β-gt* glucosyl transferase genes (Figure 1A). Using the BzCas13b system, we observed important variability across different protospacers, similarly to what was observed with LseCas13a in targeting of phiKZ, despite using the recommended 5’ and 3’ PFS (31). We observed that the presence of cytosine at the 5’ PFS end of the protospacer leads to inhibition of targeting (Figure 5), in line with previous observations with BzCas13b. Differences in expression timing, levels of the protospacer containing transcript and formation of RNA secondary structures encompassing the target sequences are possible explanations for the differences seen among protospacers of different genes and between protospacers targeting the same gene.

Finally, we found that Cas13b mediated T4 genome modification can be achieved very rapidly. First, the high efficiency of recombination within the T4 phage allowed us to perform gene deletions and insertion, as well as point mutations, with edit flanking homologous arms under 100 bps long, which greatly facilitates the construction of the plasmids for recombination. Previous studies have utilized individual flanking sequences in length greater than 250 bps for editing, exceptionally around 50 bps flanking sequence were utilized in a single case for the introduction of mutations (34). Second, despite the short edit-flanking homologous arms utilized, only 1 to 2 cycles of phage propagation were necessary to obtain sufficient recombinant phages. Third, we found that direct plating on the Cas13b and crRNA expressing strain was sufficient to select the recombinants (Figure 4). Taken together, these last two points enabled the development of a 6-hour protocol for genome editing.

In conclusion, this study presents a comprehensive view of the possibility of utilization of BzCas13b for genome engineering of T4 like bacteriophages, complementing recent studies utilizing the type VI-A system for genome editing of bacteriophages.

## MATERIALS AND METHODS

### PCR, Plasmids, strains and growth conditions

*Escherichia coli* DH10B was utilized for all experiments including genome editing and plaque assays. All *E. coli* cultures were grown in Luria-Bertani (LB) medium, incubated at 37°C with shaking at 200 rpm. Oligonucleotides containing sequences corresponding to the homologous regions for T4 genome deletion and mutation were fused together by Overlap Extension-PCR and cloned into the plasmid pBAD24 via Circular Polymerase Extension Cloning (46). In case of *yfp* insertion into the genome of T4, oligonucleotides with the homologous regions were used to amplify the *mVenusNB* gene. The amplified product was cloned into the pBAD24 plasmid via Circular Polymerase Extension Cloning. Bacteriophage targeting spacers were introduced into plasmids pC0003 (#79152), containing the Cas13a locus from *Leptotrichia Shahii*, and pBzCas13b (#89898), encoding the Cas13b from *B. zoohelcum* ATCC 43767, both obtained from Addgene. *mVenusNB* gene was derived from pEB1-mVenusNB, obtained from Addgene (plasmid #103986). Overlapping oligonucleotides with 5′ and 3′ extensions were mixed together at a final concentration of 1 μM with PCR reaction buffer, incubated at 95°C for 10 minutes and subsequently slow cooled to facilitate annealing. T4 targeting plasmids were constructed by ligating BsaI digested plasmids with the annealed spacers. All primers utilized in this study are listed in Supplementary Table 2. All polymerase chain reactions (PCRs) were performed with the respective primers using the Phusion High-Fidelity DNA Polymerase (NEB).

### Phage preparation, plaque assay, spot assay and EOP (Efficiency of plating)

To obtain phage stocks, *E. coli* DH10B culture at an OD_600_ of 0.2 was infected with phage T4 at a MOI comprised between 0.01 and 0.1, and grown at 37°C with shaking until complete lysis was observed. After centrifugation to pellet the bacterial cells, supernatants containing phage were filtered using a 0.2 µm sterile filter to eliminate remaining cells.

For plaque assay, 100μl overnight bacterial culture was mixed with preheated top agar (0.4% agar) along with the respective phage dilution and poured onto LB agar plates (containing the appropriate antibiotics, when necessary). The plates were incubated overnight at 37°C unless otherwise specified. For spot assay, 3 to 5μl of phage dilutions were spotted onto a solidified layer of 3 ml of top agar containing 100 μl of overnight bacterial culture. EOP were estimated by spot assay, by spotting serial dilutions of the T4 wild-type onto top agar mixture containing *E. coli* DH10B culture with the respective Cas13-spacer plasmid. EOP were obtained by dividing the concentration of T4 plaque forming units (PFU) on the strain of interest by the PFU concentration on DH10B.

### Genome editing - long protocol

Overnight culture of the *E. coli* DH10B recombination strain was diluted 1000-fold and incubated until an OD_600_ of 0.2 was reached. Serial dilutions of phage T4 were prepared, and 10μl of each dilution were mixed with 190μl of the recombination strain in 96-well microtiter plates and incubated at 37°C until complete lysis was observed at all dilutions, approximately between 4 to 5 hours. In the second step, similarly, overnight culture of the selection strain was diluted 1000-fold and incubated until an OD_600_ of 0.2 was reached. Supernatants obtained previously with recombination strain were serially diluted, and 10μl of each dilution were mixed with 190μl of the selection strain in 96-well microtiter plates and incubated at 37°C for 8-12 hours. Wells with clear growth retardation were screened for positive phage recombinants with PCR. Single plaques of the recombinant phage were then obtained following plaque assay on the selection strain.

### Genome editing - short protocol

Exponentially growing *E. coli* DH10B recombination strain at OD_600_ of 0.2, prepared by 100- fold dilution of overnight culture, was infected with T4 phage at an MOI of 0.1 and incubated at 37°C for 60 minutes. Serial dilutions of the supernatant were spotted on the selection strain (as described earlier) and incubated at 37°C for five hours. Single plaques were screened for the appropriate mutation by PCR.

### One-step growth curve

Phage burst sizes were measured as described earlier (47). Briefly, 10 μl of phage (10^7^ PFU/ml) were mixed with 100 μl of *E. coli* DH10B, concentrated from 1 ml of exponentially growing culture (at OD_600_= 0.2) in LB broth. The mixture was incubated at 37°C for 5 minutes, centrifuged, washed once with preheated LB. Infected cells were diluted 100-fold and 10,000- fold with preheated LB and incubated at 37°C. Samples were withdrawn at the appropriate time points and phage titers estimated by plaque assay.

### DNA extraction and sequencing

Preparations of the wild-type and mutant phage DNA were performed as described earlier. Phages were precipitated overnight using PEG (10%, final concentration) and NaCl (0.5 M, final concentration), centrifuged at 12,000 rpm for 30 minutes and resuspended in SM buffer (50 mM Tris-HCl pH7.5, 100 mM NaCl, 8 mM MgSO4). Standard Phenol-Chloroform extraction, followed by ethanol precipitation was performed to extract total phage DNA. DNA sequencing was performed by Next Generation Sequencing (INVIEW virus, Eurofins Genomics, Ebersberg, Germany). The sequencing reads were assembled using SPAdes (48) version 3.15.3 with default parameters. Mutations in T4 genomes were identified with breseq (49) version 0.36.1, using default parameters with the wild-type T4 genome as the reference sequence.

### Fluorescent *E. coli* strain construction, microscopy and Image analysis

MG1655 PRNA1::*tdCherry E. coli* strain was constructed using plasmid pNDL-32 obtained from Johan Paulsson’s lab (http:/openwetware.org/wiki/Paulsson:Strains, as described before (50), Supplementary Table 3). Cells were grown up to an OD_600_ of 0.5 in LB at 37°C, from a 200-fold dilution of overnight culture, and T4 phage was added at an MOI of 1. Immediately after, the cell and phage mixture was spread on 1% agarose + LB pad in a microscope slide, using a Thermo Scientific™ Gene Frame. Image acquisition started ∼20-25 minutes after, just as the first cell lysis occurred. We used an inverted Nikon Eclipse Ti-E microscope equipped with a KINETI sCMOS camera, a CoolLED pE-800 light source with LED wavelengths at 580 nm for tdCherry and 500 nm for YFP, a Plan APO 100x oil immersion objective (N.A. 1.45), and an Okolab temperature-controlled chamber regulated at 37°C. YFP and tdCherry exposure time and intensity were 500 ms, 5% and 200 ms, 2% respectively. Images were taken every 2 minutes for 2 hours and image analysis was performed with OUFTI© software (51), using the cytoplasmic tdCherry signal for cell segmentation. Mean pixel intensity from YFP channel, for the selected lysing cells, were extracted and plotted using Matlab Mathworks© suite.

## DATA AVAILABILITY

Data supporting the findings in this study are available within the paper and its supplementary information. Additional data can be obtained from the corresponding author upon request.

## Author contribution statement

**Yuvaraj Bhoobalan-Chitty:** Conceptualization, Investigation, Formal analysis, Supervision, Validation, Funding acquisition, Writing – original draft, Writing – review and editing.

**Mathieu Stouf:** Investigation, Formal analysis, Validation, Writing – original draft.

**Marianne De Paepe :** Conceptualization, Supervision, Validation, Funding acquisition, Writing – review and editing.

## ACKNOWLEDGEMENTS

All members of the MuSE Lab at Micalis, INRAe, Jouy-en-Josas are thanked for their suggestions. This work was supported by the Novo Nordisk Fonden Postdoctoral Fellowship in Bioscience and Basic Biomedicine Grant [NNF21OC0067491] to Y.B.-C., and ANR grant MUMI [ANR-20-CE12-0008-02] to M.D.P.

