## Supplementary File 1 for "Genetic manipulation of Bacteriophage T4 utilizing the CRISPR-Cas13b system"

### SUPPLEMENTARY FIGURES

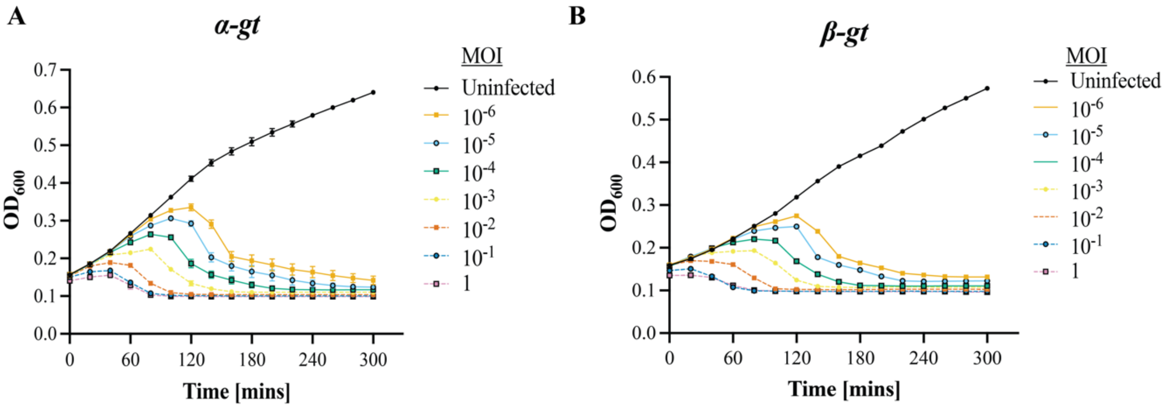


**Supplementary Figure 1. Growth curves of α-gt and β-gt recombination strains infected with wild-type T4. A**. Growth curve of the recombination strain carrying the plasmid for the deletion of alpha glucosyltransferase (α-gt) at different MOIs. **B.** Growth curve of the recombination strain carrying the plasmid for the deletion of beta glucosyltransferase (β-gt) at different MOIs.

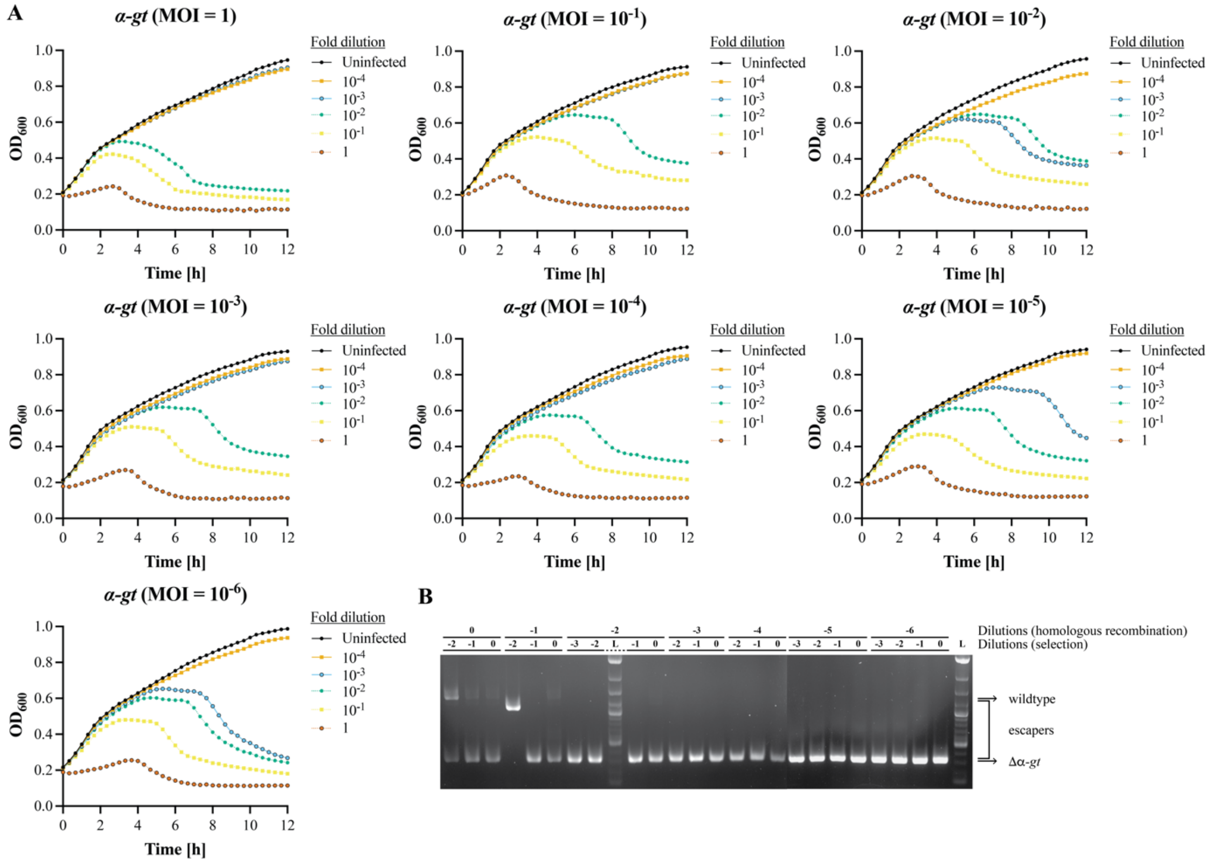


**Supplementary Figure 2. Growth curves of α-gt selection strain infected with a mixture of recombinant and wild-type T4.** A. Growth curves of the selection strain, carrying the plasmid-borne Cas13b and a spacer targeting *α-gt*, infected with serial dilutions of the phage lysates obtained from the infection of the recombination strain (Supplementary Figure 1A), containing a mixture of recombinant (Δ*α-gt*) and wild-type phages. **B.** PCR products of *α-gt* locus in phage lysates of the selection strain. Expected sizes of wild-type and deleted genes are shown by the arrows on the right. Intermediate sizes correspond to unattended deletions.

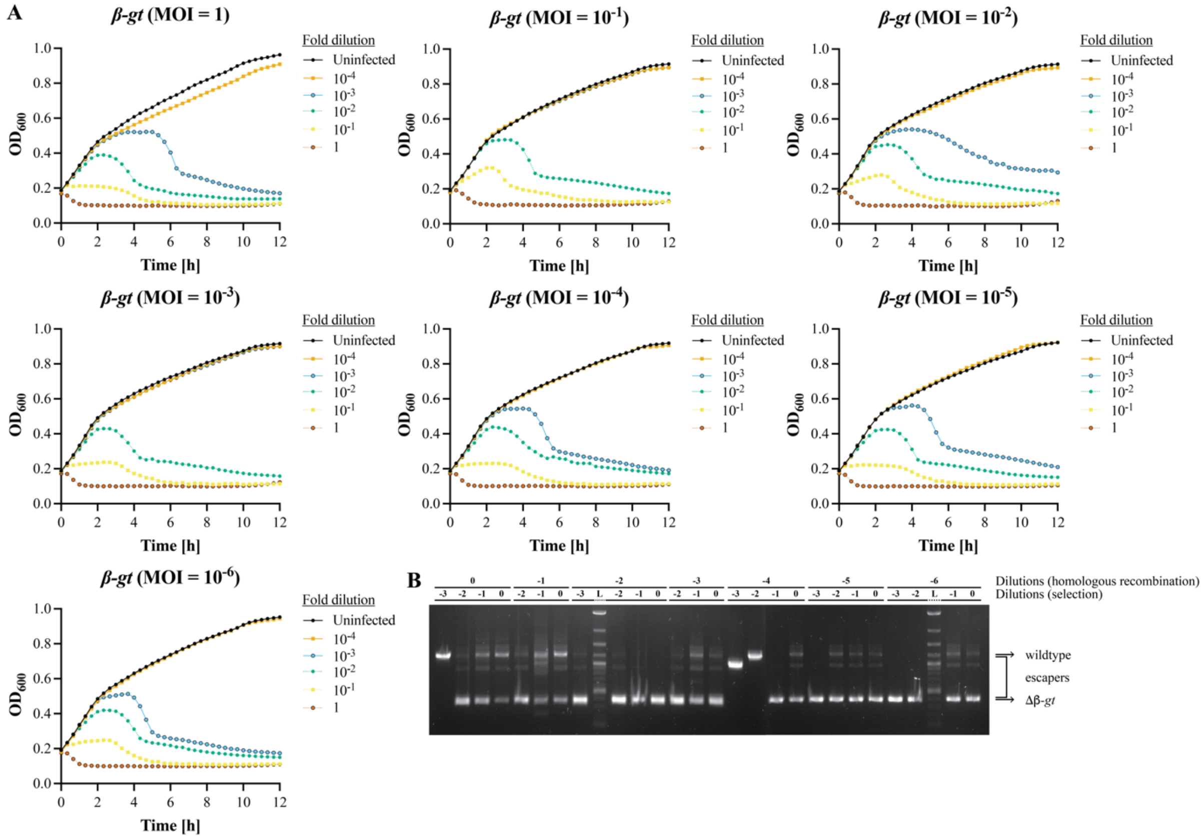


**Supplementary Figure 3. Growth curves of *β*-gt selection strain infected with a mixture of recombinant and wild-type T4.** Growth curves of the selection strain, carrying the plasmid-borne Cas13b and a spacer targeting *β-gt*, infected with dilutions of phage lysates obtained from infection of the *β-gt* recombination strain (Supplementary Figure 1A), containing a mixture of recombinant (Δ*β-gt*) and wild-type phages. **B.** PCR products of *β-gt* locus in phage lysates after growth on the selection strain. Expected sizes of wild-type and deleted genes are shown by the arrows on the right. Intermediate sizes correspond to unattended deletions.

***breseq*** version 0.36.1
mutation predictions | marginal predictions | summary statistics | genome diff | command line log

| Predicted mutations | | | | | |
|---|---|---|---|---|---|
| **evidence** | **position** | **mutation** | **annotation** | **gene** | **description** |
| MC JC | 36,721 | Δ1,065 bp | coding (29-1093/1203 nt) | *agt* ← | DNA alpha-glucosyltransferase |
| RA | 159,371 | C→T | T939I (ACT→ATT) | *T4_gp_00251* → | Long-tail fiber protein gp37 |

***breseq*** version 0.36.1
mutation predictions | marginal predictions | summary statistics | genome diff | command line log

| Predicted mutations | | | | | |
|---|---|---|---|---|---|
| **evidence** | **position** | **mutation** | **annotation** | **gene** | **description** |
| MC JC | 24,429 | Δ838 bp | | *[bgt]* | *[bgt]* |
| RA | 59,069 | T→G | H42P (CAC→CCC) | *T4_gp_00102* ← | hypothetical protein |
| RA | 84,622 | A→G | N528S (AAC→AGC) | *7* → | Baseplate wedge protein gp7 |

**Supplementary Figure 4. Mutations present in T4Δ*α-gt* (top) and T4Δ*β-gt* (bottom).** Analysis of whole genome sequencing reads with Breseq computational pipeline revealed the expected deletions, as well as one (in T4Δ*α-gt*) and two (in T4Δ*β-gt*) amino acids substitutions in the recombinant phages as compared to the wild-type T4.

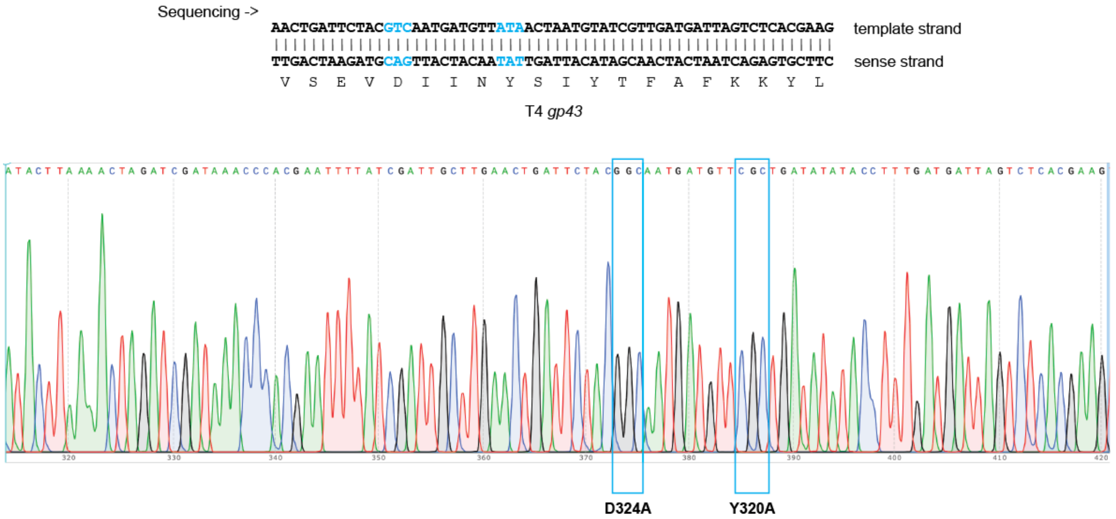


**Supplementary Figure 5. Sequence of the T4 DNA polymerase exonuclease gene region comprising the mutations introduced in T4DNApolMut01.** Top panel shows the sequence of the wild-type T4 phage with the direction of sequencing indicated. Mutated nucleotides are in blue. The bottom panel shows the sequence chromatogram corresponding to the mutant phage T4DNApolMut01.

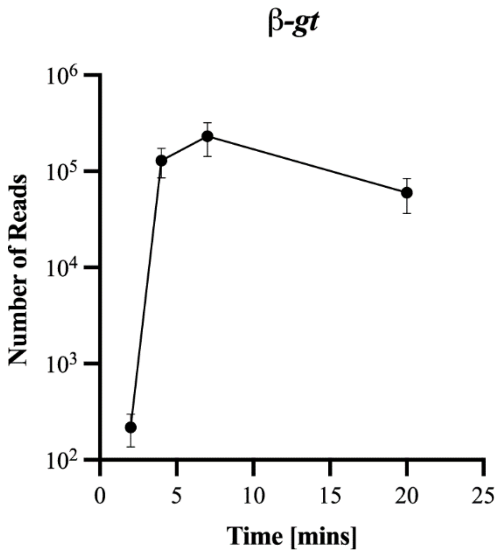


**Supplementary Figure 6.** ***β-gt*** **expression level during T4 infection cycle.** Number of *β-gt* reads plotted against time. Data derived from raw sequencing data of Wolfram-Schauerte et al ., (1). Data shown are the mean of three biological replicates, represented as mean value ± SD.

```
TATGTAGGAAAATACGCTTAATCGTTTAACATAAA--AGGAAT--AATATG   β-gt (wildtype T4)
||||||||||||||||||||||||||||||||    |   ||||||   ||||||
TATGTAGGAAAATACGCTTAATCGTTTAACAT--AGGAGGAATTCAATATG   mVenus NB
```

**Supplementary Figure 7. Upstream sequence of** ***mVenusNB*** **in comparison with that of** ***β-gt*****.** Nucleotide comparison of the upstream sequence of *β-gt* in wild-type T4 and the intended sequence, with modification to the SD sequence (highlighted in red), to be included during insertion of *mVenusNB* gene. The start codon of β-gt and mVenus NB are shown in blue.

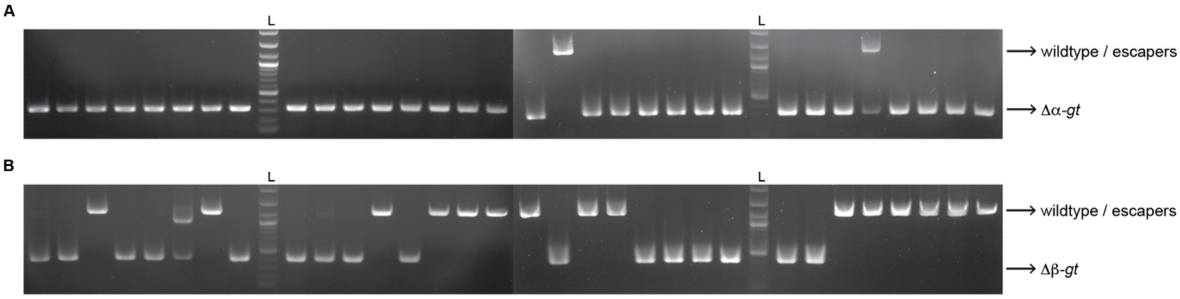


**Supplementary Figure 8. PCR analysis of single plaques of lysis obtained by the fast genome editing protocol. A**. PCR products of T4 single plaques obtained upon direct plating on the selection strain (DH10B/pCas13b-Spc(*α-gt*)), after one hour of incubation in the recombination strain (DH10B/pBAD24-HA(*α-gt*)). Arrow pointing to Δ αgt indicates the fragment size expected with the intended deletion. Arrow pointing to wild-type/escapers indicates the size expected with the wild-type gene. L: 1kb+ DNA ladder. **B.** Similar to A, but with the selection and recombination strain corresponding to *β-gt* deletion.

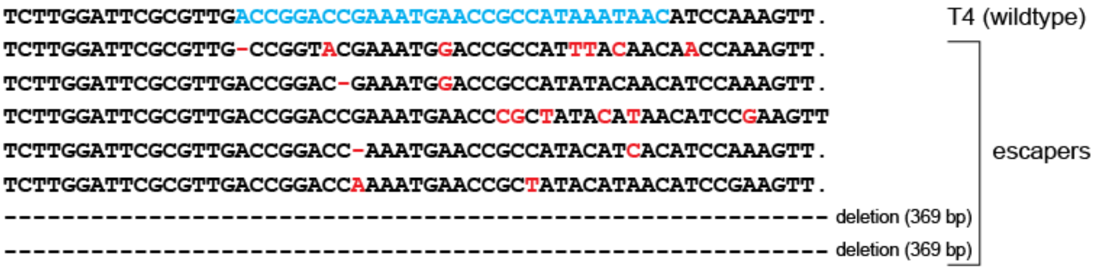


**Supplementary Figure 9. CRISPR-Cas13b targeting gives rise to T4 escape mutants.** Sequences of escape phages obtained upon targeting of T4 with a Cas13b spacer specific to *β-gt*. Two escape mutants showed deletion across the protospacer region due to microhomology mediated recombination across the sequence TTGCCAWAA, while other contain several point mutations.

**Supplementary Table 1: Cas13 spacers used in this study**

| **Type VI – subtype** | **Target gene** | **Sequence (5′ → 3′)** |
|---|---|---|
| VI-A | *α-gt* | TTACACCACAACCTTCAAGACCTCGAGCCA |
| VI-A | *β-gt* | CCATAAATTTTTGCGCAGATAAAATTGCTA |
| VI-B | *α-gt* | ATTCCATATCATTTTCATCAAACCAAATGA |
| VI-B | *β-gt* | AATAGAAAGCATCTCTTTACGCAAAACATC |
| VI-B | *gp43* (*DNApol*) | TAACTAATGTATCGTTGATGATTAGTCTCA |
| VI-B | *denA* | TATACTTCCATTGTCGTCTAATTCTAGCTC |
| VI-B | *Alc* | TCCCGTCAAAGGATTAACGATGATATAGCA |
| VI-B | *gp56* | GTTCTTCAGATAATAAAGTTTAAAGATTTC |
| VI-B | *gp43* (*DNApol*) | AACTCTGCTGAGTTTCCATACCCATGATTT |
| VI-B | *gp43* (*DNApol*) | AGGAATAACCTTATGTTCACCTTTCAATGA |
| VI-B | *res* | TCGTAATGAATGCACAACTTTTTCTAGTTC |
| VI-B | *res* | CCTCTACATATAGCTGATTTGCATATTGAA |
| VI-B | *dexA* | GAAGAGATAGAGGATCGCATTCTTCCTCTG |
| VI-B | *gp3* | CAGTACATACTAAAGCCGGGTCTGAATCTT |
| VI-B | *gp7* | GAGATAATGGATCAGAATCTATAGGTGCAT |
| VI-B | *gp61* | CCTCAGGATAAGCTTTGATGGTGATATATT |
| VI-B | *gp6* | AACCAGCATGAACCATTGACTTTCTCGTCC |
| VI-B | *gp6* | GGTTTCGCGTTTAATAGTACCCAATTCGCG |
| VI-B | *Vs. 6* | ACCACCTTCAATTTTAACTGTAGGTTGTGG |
| VI-B | *gp15* | GTCTATTTACATTACTAGTGAAAGTTTGTT |
| VI-B | *gp10* | TAATCATTAACTGTACCCTTTGGAAGATTT |
| VI-B | *gp10* | GCGGAGTGTGAGTAGAGTTAGTAGATGCTT |
| VI-B | *gp34* | GGTTAAGGTTAGTGAACCATTAACCGTCTG |
| VI-B | *hoc* | TCGCTTCAGCCGTTTCCGGGCCTCCTTCAG |
| VI-B | *rIIA* | TATTAATACCAAACAGACGATAACTTGATG |
| VI-B | *gp18* | CACAAACTCATTTCTATCAATTACTGACG |
| VI-B | *gp18* | TATCGCAAACTACACGATATTCATAAATTC |
| VI-B | *gp12* | ACGATGCATCAGGAACTCCATTTACTCCAG |
| VI-B | *gp12* | CCAACGTTGCTGGTGTAACTGCTTTGGTAT |
| VI-B | *uvsW* | GAACAATGATAAGAATTTTACCTTCATAAT |
| VI-B | *gp42* | CCAAATCTTCGGTGTTTCACCCGGAATATC |
| VI-B | *dda* | ATGTAGGAAGTGCTTTCACTTTACTAAACT |
| VI-B | *gp30.3* | TTCAACATTTTTCACCTTACACCCTTGGAG |
| VI-B | *gp23* | GTAGCATTGTAACCGTGGTCACCACCGATT |

#### Supplementary Table 2: Primers used in this study

| Plasmid construct | Oligonucleotide | Sequence (5′ → 3′) |
|---|---|---|
| pC0003-Spc(*α-gt*) | T4_*α-gt*_Cas13a_for | AAAC *TTACACCACAACCTTCAAGACCTCGAGCCA* |
| | T4_*α-gt*_Cas13a_rev | TATC *TGGCTCGAGGTCTTGAAGGTTGTGGTGTAA* |
| pC0003-Spc(*β-gt*) | T4_*β-gt*_Cas13a_for | AAAC *CCATAAATTTTTGCGCAGATAAAATTGCTA* |
| | T4_*β-gt*_Cas13a_rev | TATC *TAGCAATTTTATCTGCGCAAAAATTTATGG* |
| pCas13b-Spc(*α-gt*) | T4_*α-gt*_Cas13b_for | ACAAC *ATTCCATATCATTTTCATCAAACCAAATGA* |
| | T4_*α-gt*_Cas13b_rev | CAAC *TCATTTGGTTTGATGAAAATGATATGGAAT* G |
| pCas13b-Spc(*β-gt*) | T4_*β-gt*_Cas13b_for | ACAAC *ACCGGACCGAAATGAACCGCCATAAATAAC* |
| | T4_*β-gt*_Cas13b_rev | CAAC *GTTATTTATGGCGGTTCATTTCGGTCCGGT* G |
| pBAD24-HA(*α-gt*) | T4_*α-gt*_HA_for | ACAGCCAAGCTTGCATGCCTGTTCTTTAAAGCAGAAGCTTGAATCTTGATGCTGATACAAAAATTCATATGCTTTTTCTCGCTCACGGTCATAAAGAGCTCGGTCAGCTCGAGCCAT |
| | T4_*α-gt*_HA_rev | GAGGAATTCACCATGGTACCCGTTTTATAGAAAATAAAATATTATTTACATGATTTATTAAATGAAAAGAGGAAAACTATGCGTATTTGCATTTTTATGGCTCGAGCTGACCGAGCT |
| pBAD24-HA(*β-gt*) | T4_*β-gt*_HA_for | ACAGCCAAGCTTGCATGCCTGTTTTGTGAAATTTTTTAAATGGAAAGATACCATCCGTTGTAGTTGCTTTTTCTTACAACTTTACGAAGGTCTTCTCTGTCACCGACACTGTTCGAT |
| | T4_*β-gt*_HA_rev | GAGGAATTCACCATGGTACCCGACATAAAGGAAAGTTTAAATGCAGAAAACGAATCCTGGGTTACAGAGACTATTTCAGATTCCGACATTTACCCTATCGAACAGTGTCGGTGACAGAG |
| T4 Δ*α-gt* check | T4_*α-gt*_chk_for | ATCCAACATGCTCTAGTGAATAG |
| | T4_*α-gt*_chk_rev | GATATGTTTGGGCCTATTCGTA |
| | T4_*α-gt*_chk_in_for | TATCAATCTCACGATTACCGTA |
| | T4_*α-gt*_chk_in_rev | TACAAGTTCTCATGACCACAA |
| T4 Δ*β-gt* check | T4_*β-gt*_chk_for | TCCTAACATTTATTCCACCGGT |
| | T4_*β-gt*_chk_rev | CCTACATGTGTATTCTCGTCAT |
| | T4_*β-gt*_chk_in_for | CTCATTGACTCTATCAATGAGTTC |
| | T4_*β-gt*_chk_in_rev | TTGTACACTGAAGAAGAGCTAT |
| pCas13b-Spc(*pol*Y320) | T4_*DNApol* Y320_for | ACAAC *TAACTAATGTATCGTTGATGATTAGTCTCA* |
| | T4_*DNApol* Y320_rev | CAAC *TGAGACTAATCATCAACGATACATTAGTTA* G |
| pBAD24-HA(*pol*Y320) | T4_*DNApol*_Y320A/D324A_HA_for | ACAGCCAAGCTTGCATGCCTGCTTAAAACTAGATCGATAAACCCACGAATTTTATCGATTGCTTGAACTGATTCTACGGCAATGATGTTCGCTGATATATACCTTTGATGATTAGTCTCA |
| | T4_*DNApol*_Y320A_HA_rev | GAGGAATTCACCATGGTACCCGTCAGTTGCTCAACATGAAACCAAAAAAGGTAAATTACCATACGACGGTCCTATTAATAAACTTCGTGAGACTAATCATCAACGATATATATCAGC |
| *DNApol* Y320AD324A mut check | T4_*DNApol*_chk_for (for sequencing) | ATTCGGAGAACAAGAATATTCATCAC |
| | T4_*DNApol*_chk_rev (for sequencing) | GATGAAGCGAATGGAAGACATCG |
| | *DNApol*Y320Amut_chk_rev | CTAATCATCAACGATATATATCAGCGA |
| pBAD24-HA(*β-gt-mVenusNB*) | *mVenusNB-β-gt*-For | TATGTAGGAAAATACGCTTAATCGTTTAACATAGGAGGAATTCAATATGAGTAAAGGAGAAGAACTTTTCAC |
| | *mVenusNB-β-gt* -Rev | AATAATAGTTCATAATTTTTATTTGTATAGTTCATCCATGCCAT |

Sequences corresponding to Cas13 spacers are italicized.

#### Supplementary Table 3: Bacterial strains used in this study

| Strain name | genotype | note |
|---|---|---|
| DH10B | $F^-$, mcrA, Δ(*mrr-hsd*RMS-*mcr*BC),Φ80d*lac*ZΔM15Δ*lac*X74, *end*A1, *rec*A1, *deo*R, Δ(*ara*,*leu*)7697, *ara*D139, *gal*U, *gal*K, *nup*G, *rps*L, $\lambda^-$ | (2) |
| MS102 | *MG1655 [rph-1, λ-,* pRNA1::tdCherry) | This study |

#### Supplementary Table 4: Phage strains used in this study

| Strain name | genotype | note |
|---|---|---|
| T4Δ*α-gt* | Δ*α-gt* | This study |
| T4Δ*β-gt* | Δ*β-gt* | This study |
| T4DNApolMut01 | *gp43Y320AD324A* | This study |
| T4Δ*β-gt*∇*mVenus* | Δ*β-gt* ∇*mVenus* | This study |
